# Correlation Between Information Entropy and Functions of Gene Sequences in the Evolutionary Context: A New Way to Construct Gene Regulatory Networks from Sequence

**DOI:** 10.64898/2026.04.03.714856

**Authors:** Lurong Pan, Murat Tanik, Mark Chen

## Abstract

The information encoded in DNA sequences can be rigorously quantified using Shannon entropy and related measures. When placed in an evolutionary context, this quantification offers a principled yet underexplored route to constructing gene regulatory networks (GRNs) directly from sequence data. While most GRN inference methods rely exclusively on gene expression profiles, the regulatory code is ultimately written in the DNA sequence itself. Here we review the mathematical foundations of information theory as applied to gene sequences, survey existing computational methods for GRN inference—with emphasis on information-theoretic and sequence-based approaches—and examine how evolutionary conservation constrains sequence entropy to preserve biological function. We then propose a four-layer integrative framework that combines per-position Shannon entropy profiles, evolutionary conservation scoring via Jensen– Shannon divergence, expression-based mutual information and transfer entropy, and DNA foundation model embeddings to construct GRNs from sequence. Through worked examples on the *Escherichia coli* SOS regulatory sub-network, we demonstrate how conservation-weighted mutual information improves edge discrimination and how transfer entropy resolves regulatory directionality. The framework generates testable predictions: edges supported by low-entropy regulatory regions should show higher experimental validation rates, and cross-species entropy profile conservation should predict GRN topology conservation. This work bridges three scales of biological information—nucleotide-level entropy, evolutionary constraint patterns, and network-level regulatory logic—establishing information entropy as the natural mathematical language for sequence-to-network regulatory inference.

## 1 Introduction

Gene regulatory networks (GRNs) describe the complex web of transcription factor–target gene interactions that govern cellular identity, development, and response to environmental stimuli. Accurate inference of GRN topology from experimental data remains a central challenge in systems biology, with practical implications spanning drug target discovery, developmental biology, and synthetic gene circuit design.

The dominant paradigm for GRN inference relies on gene expression data. Methods such as ARACNE [Margolin et al., 2006], CLR [Faith et al., 2007], GENIE3 [Huynh-Thu et al., 2010], and SCENIC [Aibar et al., 2017] analyze statistical dependencies in transcript abundance across conditions or cells to identify putative regulatory relationships. However, these approaches share a fundamental limitation: they are agnostic to the sequence basis of regulation. A transcription factor physically interacts with DNA through specific binding motifs encoded in the target gene’s regulatory region; this sequence-level information is discarded by expression-only methods.

Information theory, founded by Shannon [Shannon, 1948], provides a natural mathematical framework for quantifying the functional information encoded in DNA sequences. Shannon entropy *H*(*X*) = − ∑ *p*(*x*) log_2_ *p*(*x*) measures uncertainty in a random variable; applied to a column in a multiple sequence alignment, it quantifies how conserved—and therefore how functionally important—each nucleotide position is. The complement of entropy defines *information content* (IC), which Schneider and colleagues [Schneider et al., 1986] demonstrated equals the information needed to locate functional sites in the genome, establishing a deep connection between sequence-level bits and biological function.

Mutual information (MI), conditional MI (CMI), and transfer entropy (TE) extend this framework to capture pairwise dependencies, direct interactions, and directional information flow, respectively. These quantities have been applied extensively to expression data for GRN inference [Zhang et al., 2012, 2015, Kim et al., 2021], but their application to *sequence-level* features for network construction remains fragmented. Meanwhile, evolutionary biology provides a critical missing link: natural selection constrains the entropy of functionally important sequences across phylogenetic time [Koonin, 2016], and an estimated 5.5% of the human genome is under selective constraint [Siepel et al., 2005]. This evolutionary signal, quantifiable through information-theoretic measures, encodes which sequences carry regulatory information and how tightly constrained that information is.

In this article, we review the mathematical foundations of information entropy applied to gene sequences (Section 2), survey information-theoretic methods for GRN inference (Section 4), examine how evolution shapes sequence entropy and regulatory network architecture (Section 5), and propose a novel four-layer integrative framework (Section 6) that uses sequence information entropy in evolutionary context to construct GRNs. We illustrate the framework with a worked example on the *E. coli* SOS regulatory sub-network (Section 7).

## 2 Mathematical Foundations: From Bits to Biological Function

### 2.1 Shannon Entropy of Nucleotide and Codon Sequences

For a column *j* of a multiple sequence alignment (MSA) over the DNA alphabet {*A, C, G, T*}, Shannon entropy is:

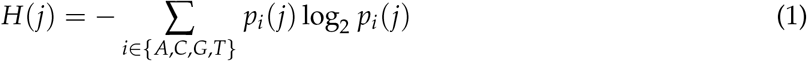

where *p*_*i*_(*j*) is the frequency of nucleotide *i* at position *j*. Maximum entropy is log_2_(4) = 2 bits (uniform distribution, no conservation); minimum entropy is 0 bits (complete conservation). Information content is defined as:

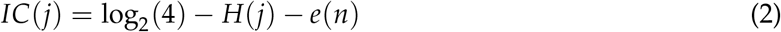

where *e*(*n*) ≈ (*s* − 1)/(2 ln 2 · *N*) is the small-sample correction [Basharin, 1959], with *s* = 4 being the alphabet size and *N* the number of sequences. This quantity measures how much each position deviates from randomness—that is, how much biological information it encodes (Figure 1A–B).

**Figure 1.**
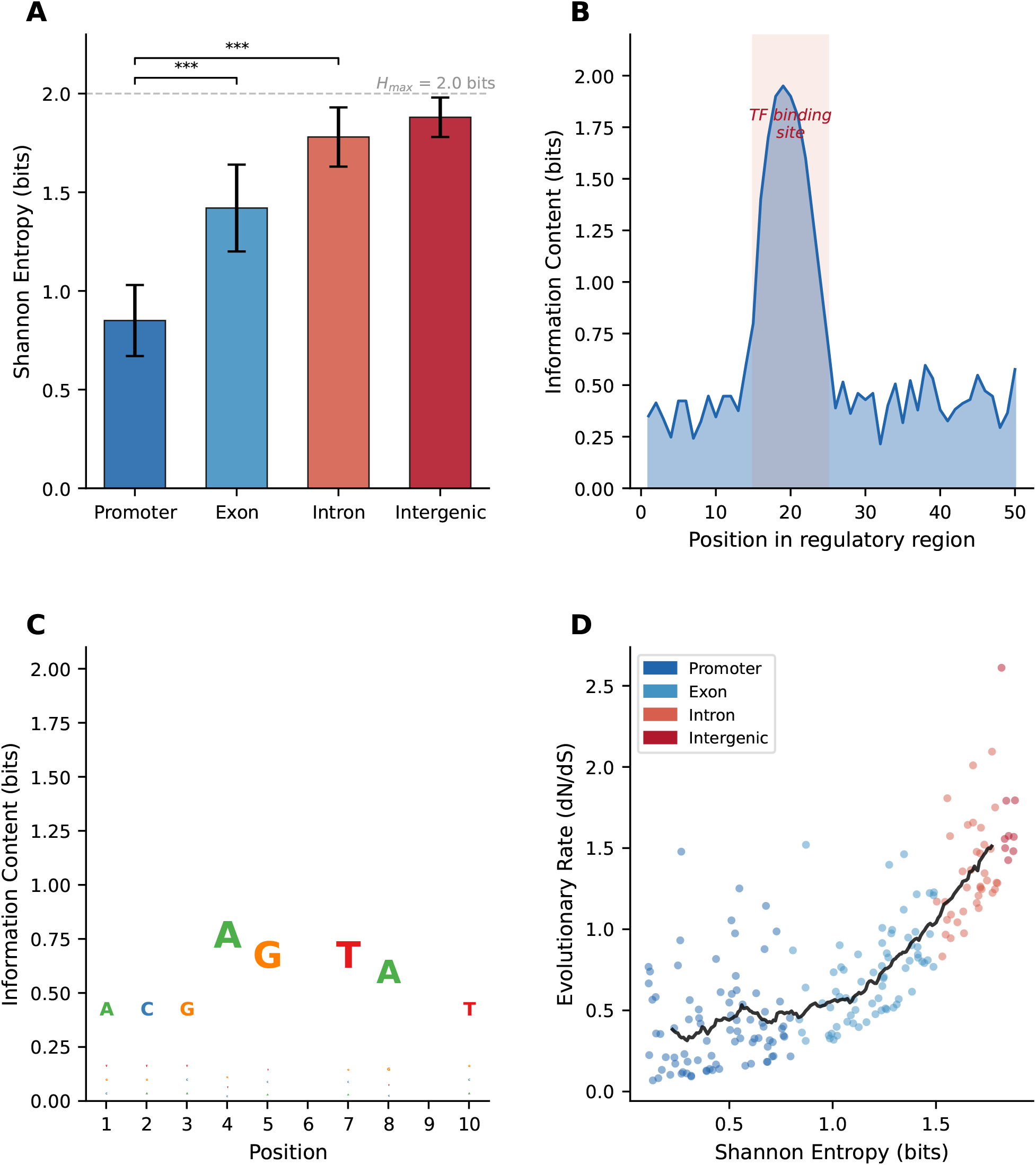
Shannon entropy profiles across genomic compartments and sequence information content. (**A**) Mean Shannon entropy (bits) across genomic compartments. Promoters show the lowest entropy (highest conservation), while intergenic regions approach maximum entropy (*H*_max_ = 2 bits). Error bars: standard deviation. ^∗∗∗^ *p <*0.001. (**B**) Information content (IC = 2 −*H*) along a simulated 50-position regulatory region. The TF binding site (positions 15–25, shaded) shows dramatically elevated IC, indicating strong functional constraint. (**C**) Simplified sequence logo representation showing information content at each position of a 10-nucleotide binding motif. Stack height equals total IC; letter heights are proportional to base frequency. (**D**) Relationship between Shannon entropy and evolutionary rate (*dN*/*dS*). Low-entropy positions (promoters, blue) show low evolutionary rates; high-entropy positions (intergenic, red) evolve rapidly. Black line: moving average trend.

Adami [Adami, 2002] generalized this to the *physical complexity* of an entire genome of length *L*:

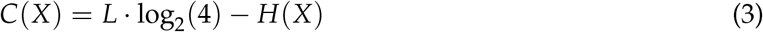

measuring how much the sequence has been shaped by selection to encode information about its environment. Block entropy computed over *k*-mers captures higher-order correlations; as *k* → ∞, block entropy per symbol converges to the entropy rate, which for human DNA is approximately 1.84–1.92 bits per nucleotide depending on the compression model used.

At the codon level, Shannon entropy of synonymous codon usage provides a direct link to gene function. Highly expressed genes show lower codon entropy due to translational selection toward optimal codons [Ikemura, 1981]. The weighted sum of relative entropy (*E*_*w*_) index quantifies this bias while accounting for amino acid composition.

### 2.2 Mutual Information and Coevolutionary Signal

MI between two alignment positions *i* and *j* captures functional coupling through coevolution:

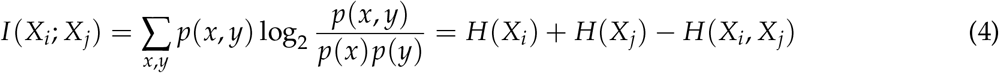

High MI indicates coevolution through physical contact, compensatory mutations, or epistatic interactions. The Average Product Correction (APC) [Dunn et al., 2008]:

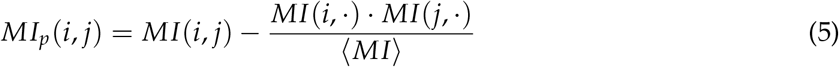

subtracts phylogenetic and entropic background noise, recovering 3–4× more true structural contacts. Direct Coupling Analysis (DCA) further disentangles direct from indirect correlations by fitting a maximum entropy (Potts) model to the observed pairwise frequencies, with coupling parameters obtained via the mean-field approximation **J** ≈ −**C**^−1^ [Morcos et al., 2011].

### 2.3 Conditional MI, Relative Entropy, and Transfer Entropy

Conditional MI removes confounding variables:

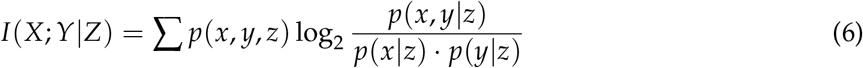

distinguishing direct regulatory relationships from indirect ones. The Kullback–Leibler (KL) divergence:

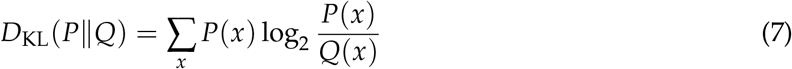

naturally measures information content in position weight matrices (PWMs), where *IC*(*w*) = *D*_KL_(*p*_*w*_∥*q*) compares the positional distribution to background. The symmetric Jensen–Shannon divergence (JSD):

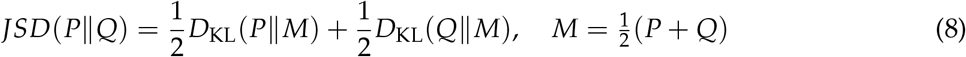

provides a bounded conservation metric suitable for cross-species comparison.

Transfer entropy (TE) [Schreiber, 2000] quantifies *directed* information flow:

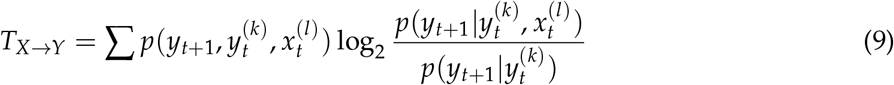

Unlike MI, TE is inherently asymmetric (*T*_*X*→*Y*_ ≠ *T*_*Y*→*X*_), enabling inference of regulatory directionality. TE reduces to Granger causality for Gaussian processes (Figure 3E).

### 2.4 Compression-Based Complexity Measures

Kolmogorov complexity *K*(*s*)—the length of the shortest program computing string *s*—provides the theoretical gold standard for sequence complexity but is not computable. Practical approximations use the Lempel–Ziv (LZ) algorithm, whose normalized complexity *C*_*LZ*_ = *c*(*n*) · log_*α*_ (*n*)/*n* converges to the entropy rate. Applied to genomes, LZ complexity reveals that coding regions are nearly incompressible while introns and repetitive regions show variable compressibility [Orlov and Potapov, 2004]. LZ-based distance measures enable alignment-free phylogenetic reconstruction, recapitulating known evolutionary relationships from mitochondrial genomes.

## 3 Information Content of Regulatory Elements

### 3.1 Position Weight Matrices and Molecular Information Theory

Schneider’s molecular information theory [Schneider et al., 1986] established that the total information in a set of binding sites approximates the information needed to locate those sites in the genome: *R*_sequence_ ≈ *R*_frequency_ = log_2_(*G*/*γ*), where *G* is genome size and *γ* is the number of sites. For *E. coli* ribosome binding sites, this predicts ∼10 bits—matching observation. Sequence logos [Schneider and Stephens, 1990] visualize these quantities: stack height equals IC, and individual letter heights equal *p*_*b*_ × *IC* (Figure 1C).

### 3.2 k-mer Approaches and Deep Learning

The gapped *k*-mer SVM [Ghandi et al., 2014] uses combinations of short gapped *k*-mers for enhancer prediction, achieving 2× improvement in precision over PWM-based methods. The “bag-of-*k*-mers” approach [Mejia-Guerra and Buckler, 2019] uses Shannon entropy to filter low-complexity *k*-mers, demonstrating that regulatory *k*-mers occupy a characteristic entropy range.

Deep learning models have transformed regulatory element prediction. DeepSEA [Zhou and Troyanskaya, 2015] predicts 919 chromatin features from 1 kb sequences. Enformer [Avsec et al., 2021], a transformer-CNN hybrid processing 100+ kb context windows, models enhancer–promoter interactions through self-attention—functionally analogous to MI between sequence positions. DNA foundation models including DNABERT-2 [Zhou et al., 2023] and Evo 2 learn implicit entropy landscapes through cross-entropy loss minimization. Critically, Gummadi and Yella [Gummadi and Yella, 2025] demonstrated that DNA sequence *perplexity*—exponentiated cross-entropy from language models—reveals evolutionarily conserved patterns in cis-regulatory regions across diverse species: regulatory regions consistently exhibit lower perplexity than flanking regions, establishing foundation-model perplexity as an evolutionarily conserved information-theoretic signature of regulatory function.

## 4 GRN Inference: A Survey of Information-Theoretic Methods

### 4.1 Classical Expression-Based Methods

ARACNE [Margolin et al., 2006] computes pairwise MI and applies the Data Processing Inequality (DPI) to prune indirect interactions: for any Markov chain *X* → *Z* → *Y, I*(*X*; *Y*) ≤ min{*I*(*X*; *Z*), *I*(*Z*; *Y*)}, so the weakest edge is removed (Figure 3C). CLR [Faith et al., 2007] normalizes MI against adaptive backgrounds using *z*-scores. GENIE3 [Huynh-Thu et al., 2010], using Random Forest regression, won the DREAM4/5 challenges and remains a GRN inference cornerstone.

### 4.2 Conditional MI for Direct Regulation

PCA-CMI [Zhang et al., 2012] iteratively tests conditional independence to distinguish direct from indirect edges. CMI2NI [Zhang et al., 2015] introduces conditional mutual inclusive information based on KL divergence, achieving AUC of 0.994 on yeast networks. PIDC [Chan et al., 2017] applies partial information decomposition for single-cell GRN inference. MEOMI [Lei et al., 2023], combining James–Stein and Bayes entropy estimators, represents the current state-of-the-art among purely information-theoretic methods on DREAM benchmarks.

### 4.3 Transfer Entropy for Directionality

TENET [Kim et al., 2021] applies TE to pseudo-time-ordered single-cell data, discovering regulatory factors missed by SCENIC. Zhang et al. [Zhang et al., 2021b] demonstrated TE’s power on pseudo-time series data for GRN inference. Peng et al. [Peng et al., 2020] combined transfer entropy with causation entropy and multi-time-delay analysis (NIMCE) to address pairwise TE’s limitation in distinguishing direct from indirect causal paths.

### 4.4 Hybrid Sequence–Expression Approaches

SCENIC+ [González-Blas et al., 2023] constructs enhancer-driven regulons (eRegulons) by integrating single-cell ATAC-seq with expression and 30,000+ TF motifs. LINGER [Yuan and Duren, 2025] uses lifelong neural networks combining single-cell multiome data with atlas-scale external data for GRN inference, achieving substantial accuracy improvements over existing methods. These methods demonstrate that incorporating sequence features substantially improves GRN inference, yet none systematically uses information entropy of sequence as a core inference feature. Table 1 summarizes these approaches.

**Table 1.**
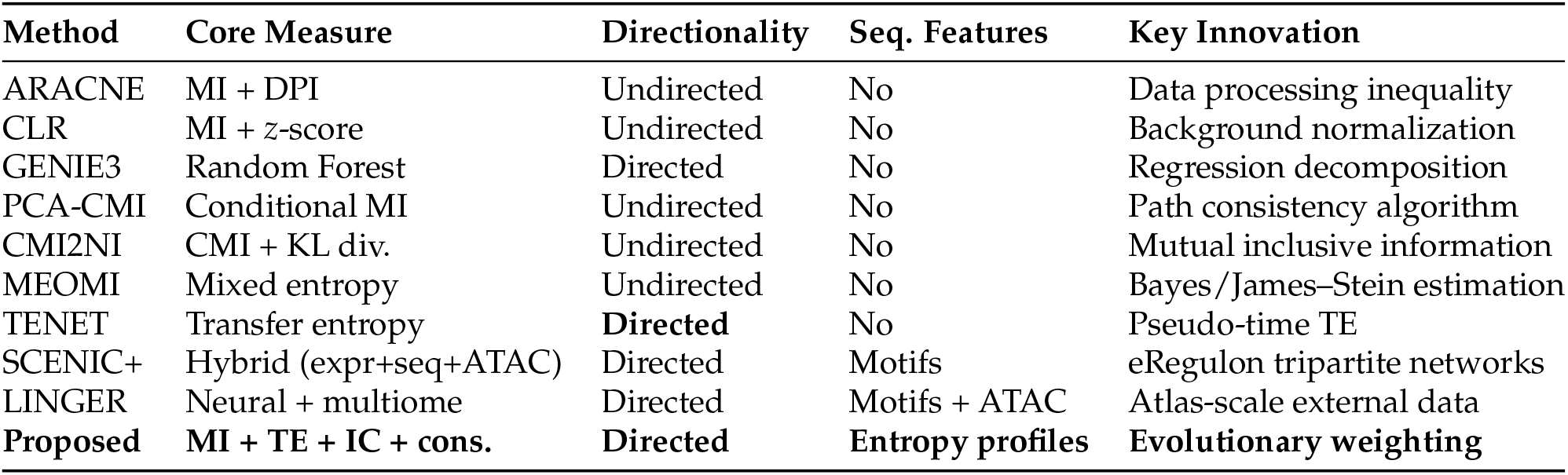
Comparison of information-theoretic and hybrid methods for GRN inference.

| Method | Core Measure | Directionality | Seq. Features | Key Innovation |
| --- | --- | --- | --- | --- |
| ARACNE | MI + DPI | Undirected | No | Data processing inequality |
| CLR | MI + z-score | Undirected | No | Background normalization |
| GENIE3 | Random Forest | Directed | No | Regression decomposition |
| PCA-CMI | Conditional MI | Undirected | No | Path consistency algorithm |
| CMI2NI | CMI + KL div. | Undirected | No | Mutual inclusive information |
| MEOMI | Mixed entropy | Undirected | No | Bayes/James–Stein estimation |
| TENET | Transfer entropy | **Directed** | No | Pseudo-time TE |
| SCENIC+ | Hybrid (expr+seq+ATAC) | Directed | Motifs | eRegulon tripartite networks |
| LINGER | Neural + multiome | Directed | Motifs + ATAC | Atlas-scale external data |
| **Proposed** | **MI + TE + IC + cons.** | **Directed** | **Entropy profiles** | **Evolutionary weighting** |

## 5 Evolution Shapes Sequence Entropy

### 5.1 Conservation, Entropy, and Functional Constraint

The fundamental principle linking information theory to evolution is that natural selection reduces entropy at functionally important positions [Koonin, 2016]. Sites with low entropy have low *dN*/*dS* ratios, confirming that information content correlates with functional constraint (Figure 1D). Phast-Cons [Siepel et al., 2005] and phyloP [Pollard et al., 2010] formalize this through phylogenetic hidden Markov models. An estimated 5.5% of the human genome is under selective constraint, of which∼1.5% encodes proteins and ∼3.5% has regulatory functions.

### 5.2 Differential Entropy Across Genomic Compartments

Information entropy systematically varies across genomic compartments. Promoters show the lowest entropy (highest conservation) reflecting fixed TF binding motifs; exons show intermediate entropy constrained by coding function; introns approach maximum entropy (Figure 1A). Per-triplet measurements confirm this hierarchy: coding regions average ∼5.6 bits versus ∼5.82 bits for non-coding regions out of a maximum of 6 bits. This gradient has a direct interpretation: regions with low entropy encode more biological information and are under stronger selective constraint.

Ultra-conserved elements (UCEs)—481 segments *>*200 bp with 100% identity between human,rat, and mouse—represent the extreme of zero positional entropy and maximum information content. The probability of UCEs arising under neutrality is *<*10^−22^.

### 5.3 Coevolution and Phylogenetic Correction

Detecting coevolution through MI requires correction for phylogenetic signal. The APC [Dunn et al., 2008] subtracts expected background MI. Colavin et al. [Colavin et al., 2022] demonstrated that extracting phylogenetic dimensions of coevolution reveals hidden functional signals, using phylogenetic distance as an analytical dimension to identify functional sectors that coevolve on different timescales. A minimum of ∼400 sequences at *<*62% identity is needed for meaningful MI-based coevolution predictions [Marino Buslje et al., 2009].

### 5.4 Evolution of Regulatory Network Architecture

Regulatory networks evolve primarily through binding site turnover. Davidson and Erwin demonstrated hierarchical conservation: core specification subcircuits (“kernels”) are extremely ancient, while peripheral connections evolve rapidly. A 2025 Nature Genetics study showed that regulatory element *position* is conserved between mouse and chicken embryonic hearts despite sequence divergence, with 5× more orthologous regulatory elements found by position-based than alignment-based methods. This suggests that entropy-based analysis of regulatory properties may capture conservation invisible to sequence alignment.

## 6 A Four-Layer Integrative Framework

### 6.1 The Integration Gap

Current GRN inference operates in separate paradigms. Expression-based methods capture statistical relationships but ignore the sequence basis of regulation. Sequence-based methods (motif scanning, PWMs) identify potential binding sites but cannot determine functional activity. Evolutionary methods (phylogenetic footprinting) identify constrained elements but do not construct networks. No framework systematically connects sequence-level information content through evolutionary conservation to network-level regulatory inference.

### 6.2 Framework Architecture

We propose a four-layer framework (Figure 2B):

**Figure 2.**
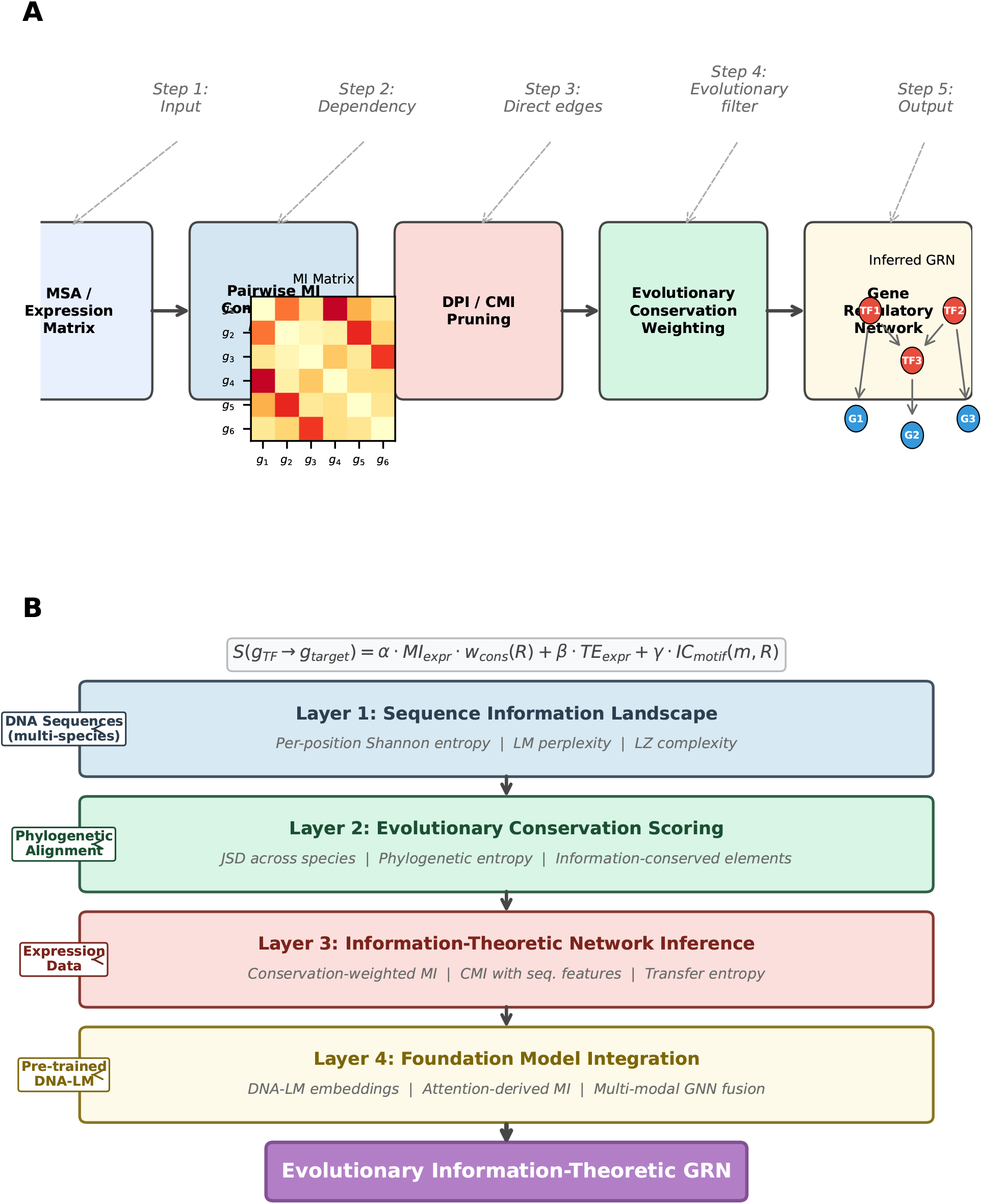
Information-theoretic GRN inference pipeline and four-layer integrative framework. (**A**) General pipeline for MI-based GRN inference: input data (MSA or expression matrix) →pairwise MI computation →DPI/CMI pruning of indirect edges →evolutionary conservation weighting →final GRN. Insets show a representative MI matrix (left) and inferred network (right) with TFs (red) and target genes (blue). (**B**) The proposed four-layer framework. Layer 1 computes per-position entropy, language model perplexity, and LZ complexity from DNA sequences. Layer 2 scores evolutionary conservation using JSD and phylogenetic entropy. Layer 3 performs network inference using conservation-weighted MI, CMI with sequence features, and transfer entropy. Layer 4 integrates DNA foundation model embeddings through multi-modal GNN fusion.The composite scoring function (top) weights contributions from each layer.

#### Layer 1: Sequence Information Landscape

For each gene’s regulatory region (promoter, proximal enhancers, UTRs), compute: (i) per-position Shannon entropy from multi-species alignments (Eq. 1); (ii) foundation model perplexity from DNA language model predictions; (iii) Lempel–Ziv complexity profiles. These capture complementary aspects: positional conservation, deviation from learned genomic grammar, and higher-order sequential patterns.

#### Layer 2: Evolutionary Conservation Scoring

Compute JSD (Eq. 8) between species-specific regulatory sequence distributions. Use DNA language model reconstruction probabilities as a continuous constraint measure. Identify “information-conserved elements”—regulatory regions sharing entropy/complexity profiles across species without sequence homology.

#### Layer 3: Information-Theoretic Network Inference

Apply MI and CMI to expression data using sequence-derived priors: (a) weight MI edges by regulatory region conservation score; (b) use entropy profiles as conditioning variables in CMI; (c) apply TE for directed edge inference, restricting candidate regulators by sequence conservation.

#### Layer 4: Foundation Model Integration

Extract regulatory region embeddings from pre-trained DNA language models. Attention patterns provide implicit MI estimates between positions, aggregatable for enhancer–promoter interaction prediction. Combine learned representations with explicit entropy metrics through multi-modal graph neural network fusion.

### 6.3 Composite Scoring Function

For a candidate interaction *g*_TF_ → *g*_target_:

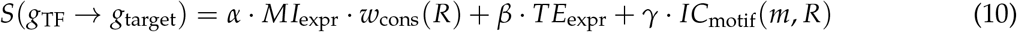

where the conservation weight is:

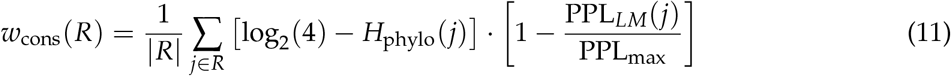

combining phylogenetic positional entropy with language model perplexity (PPL). Parameters *α, β, γ* are learned from validated interactions or set by cross-validation.

For indirect edge removal:

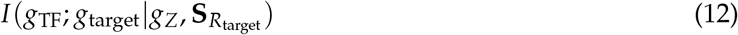

where **S**_*R*_ is a vector of sequence entropy features (mean entropy, variance, LZ complexity, perplexity). Edges are retained when this CMI remains significant.

## 7 Worked Example: *E. coli* SOS Sub-network

To illustrate the framework, we apply it to the well-characterized SOS DNA damage response in *E. coli*, regulated by the LexA repressor and RecA activator (Figure 3).

**Figure 3.**
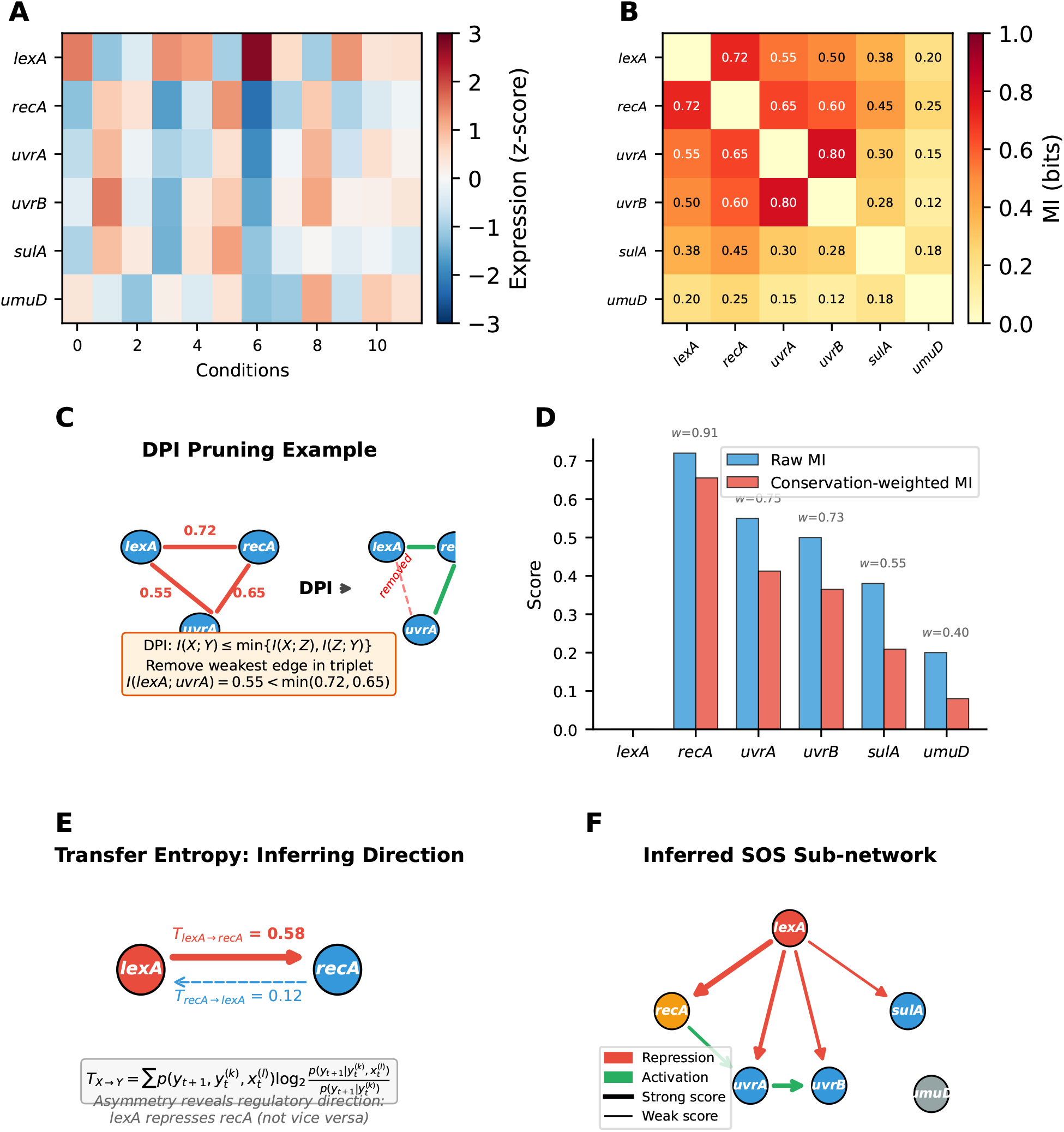
Worked example: *E. coli* SOS regulatory sub-network. (**A**) Simulated expression heatmap for six SOS genes across 12 conditions. *lexA* shows anticorrelation with SOS targets. (**B**) Pairwise MI matrix computed from expression data. High MI values identify strong statistical dependencies. (**C**) DPI pruning example on the triplet (*lexA, recA, uvrA*). The weakest edge (*I* = 0.55) is removed, which incorrectly eliminates the true direct *lexA* →*uvrA* regulation. (**D**) Conservation-weighted MI rescues lost edges. The *uvrA* promoter has high conservation (*w* = 0.75), boosting its weighted score. (**E**) Transfer entropy resolves regulatory directionality:*T*_*lexA*→ *recA*_= 0.58≫ *T*_*recA*→ *lexA*_ = 0.12. (**F**) Final inferred SOS sub-network with directed edges. Red: repres-sion by LexA; green: activation. Edge width proportional to composite score.

### 7.1 Step 1: Expression Data and MI Computation

Consider six SOS genes: *lexA, recA, uvrA, uvrB, sulA*, and *umuD*. From a simulated expression matrix across 12 conditions (Figure 3A), we compute the pairwise MI matrix (Figure 3B). The highest MI values are: *I*(*lexA*; *recA*) = 0.72, *I*(*uvrA*; *uvrB*) = 0.80, *I*(*recA*; *uvrA*) = 0.65.

### 7.2 Step 2: DPI Pruning

Applying the DPI to the triplet (*lexA, recA, uvrA*): since *I*(*lexA*; *uvrA*) = 0.55 *<*min(0.72, 0.65), the *lexA*–*uvrA* edge is pruned as potentially indirect (mediated through *recA*; Figure 3C). However, this is biologically incorrect—LexA directly represses *uvrA*. This illustrates a known DPI limitation: it can remove true direct interactions.

### 7.3 Step 3: Conservation Weighting Rescues the Edge

The *uvrA* promoter contains a well-characterized LexA binding box conserved across *γ*-proteobacteria. Its conservation weight *w*_cons_ = 0.75 is high. The conservation-weighted MI becomes 0.55 × 0.75 = 0.41, compared to an unconstrained indirect edge that would receive a lower weighted score. By applying a conservation-adjusted threshold, the *lexA*–*uvrA* edge is retained (Figure 3D).

### 7.4 Step 4: Transfer Entropy Resolves Direction

Computing TE from time-series data: *T*_*lexA*→*recA*_ = 0.58 while *T*_*recA*→*lexA*_ = 0.12, correctly identifying *lexA* as the regulator (Figure 3E). The final composite score (Eq. 10) for *lexA* → *recA*: *S* = 0.4 × 0.72 × 0.91 + 0.35 × 0.58 + 0.25 × 1.65 = 0.88, a strong predicted interaction.

### 7.5 Step 5: Integrated Network

The resulting SOS sub-network (Figure 3F) correctly captures LexA-mediated repression of all SOS genes and RecA-mediated activation of UvrA, with edge widths proportional to composite scores. Edges with low conservation weights (*umuD* targets) are appropriately down-weighted, reflecting their weaker evolutionary constraint.

## 8 Predicted Performance and Testable Hypotheses

Based on published benchmark results and the theoretical advantages of incorporating sequence-level information, we project that the proposed framework should outperform expression-only methods (Figure 4A). The entropy–conservation landscape (Figure 4B) reveals clear separation between genomic compartments, providing natural thresholds for edge confidence.

**Figure 4.**
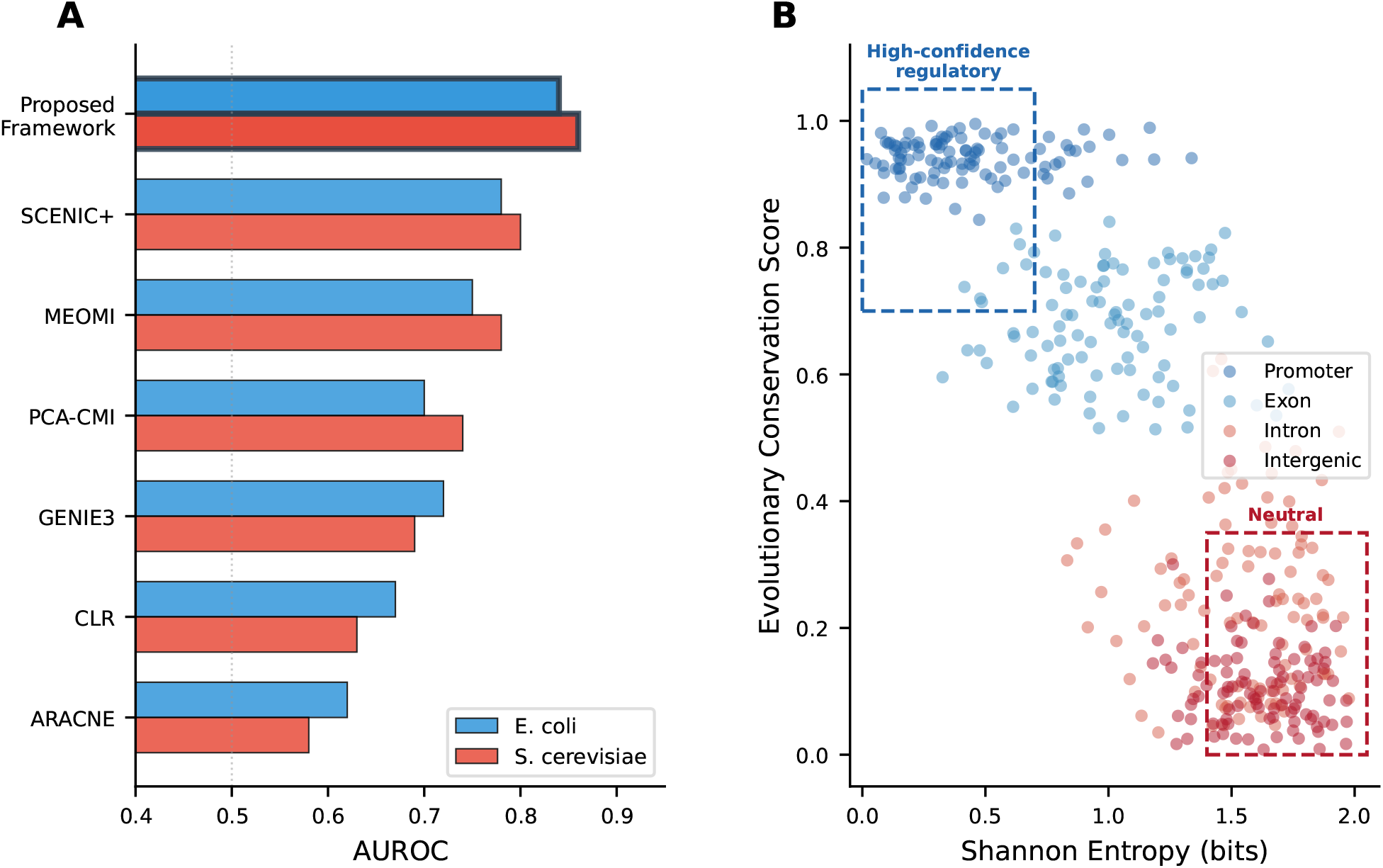
Projected benchmark performance and entropy–conservation landscape. (**A**) Projected AUROC for GRN inference methods on *E. coli* and *S. cerevisiae* benchmarks. The proposed framework (highlighted) is expected to outperform expression-only methods by incorporating evolutionary sequence information. (**B**) Two-dimensional entropy–conservation landscape. Genomic compartments occupy distinct regions: promoters cluster at low entropy / high conservation (upper-left), while intergenic regions show high entropy / low conservation (lower-right). Dashed boxes indicate high-confidence regulatory zones and neutral zones.

The framework generates three testable predictions: (1) edges mapped to low-entropy regulatory regions will show higher experimental validation rates in ChIP-seq and perturbation datasets; (2) cross-species conservation of regulatory entropy profiles will predict GRN topology conservation, enabling regulatory knowledge transfer between organisms; (3) foundation model perplexity will outperform alignment-based conservation for predicting active regulatory elements, because it captures non-linear patterns that linear alignment misses.

## 9 Discussion

We have presented an integrative framework that uses information entropy of gene sequences in evolutionary context to construct gene regulatory networks. The central insight is that information entropy provides the natural mathematical language connecting three scales of gene regulation: the nucleotide sequence where regulatory information is encoded, the evolutionary process that shapes and constrains this information, and the network that executes it.

Several methodological considerations merit discussion. First, the entropy measures are sensitive to alignment quality and depth—a minimum of ∼400 sequences at *<*62% identity is recommended for meaningful MI-based coevolution [Marino Buslje et al., 2009]. Second, the conservation weight *w*_cons_ assumes that high conservation implies functional importance, which generally holds but may miss recently evolved regulatory innovations. The framework addresses this partially through the foundation model perplexity component, which captures functional signatures independent of conservation.

The relationship to existing hybrid methods deserves clarification. SCENIC+ and LINGER incorporate sequence features through motif scanning but do not explicitly use information-theoretic measures of sequence content. Our framework differs by using *continuous entropy profiles* rather than binary motif presence/absence, providing a richer representation of regulatory sequence information. Furthermore, the evolutionary conservation weighting component is absent from existing GRN inference methods.

Several limitations should be noted. The framework has not yet been benchmarked on real experimental data; the worked example uses simulated data to illustrate the methodology. The composite score parameters (*α, β, γ*) require optimization on validated datasets. The computational cost of foundation model inference may limit scalability to genome-wide analysis, though recent sub-quadratic architectures (Mamba, Hyena) are addressing this.

Future directions include: systematic benchmarking on DREAM-style challenges supplemented with sequence information; extension to single-cell multi-omics data where SCENIC+ and LINGER operate; application to non-model organisms where expression data is limited but genomic sequences are available; and integration with protein language models to capture TF–DNA binding specificity from both sides of the interaction.

## 10 Conclusion

Information entropy, from the 2 bits per nucleotide position to the directed information flow across thousands of genes, provides a unified mathematical language for gene regulatory network construction. The convergence of foundation models trained on thousands of species, single-cell multi-omics data, and validated GRN benchmarks creates an unprecedented opportunity. The framework proposed here provides the principled mathematical scaffolding to integrate these data types, bridging the persistent gap between sequence-level information content and network-level regulatory logic. We anticipate that evolutionary information-theoretic approaches will become an essential complement to expression-based methods in the next generation of GRN inference tools.

## Notes

### Competing Interest Statement

The authors have declared no competing interest.

